# Identification of protein abundance changes in biopsy-level hepatocellular carcinoma tissues using PCT-SWATH

**DOI:** 10.1101/300673

**Authors:** Yi Zhu, Jiang Zhu, Cong Lu, Ping Sun, Wei Xie, Qiushi Zhang, Liang Yue, Tiansheng Zhu, Guan Ruan, Ruedi Aebersold, Shi’ang Huang, Tiannan Guo

## Abstract

In this study, we optimized the pressure-cycling technology (PCT) and SWATH mass spectrometry workflow to analyze biopsy-level tissue samples (2 mg wet weight) from 19 hepatocellular carcinoma (HCC) patients. Using OpenSWATH and pan-human spectral library, we quantified 11,787 proteotypic peptides from 2,579 SwissProt proteins in 76 HCC tissue samples within about 9 working days (from receiving tissue to SWATH data). The coefficient of variation (CV) of peptide yield using PCT was 32.9%, and the R^2^ of peptide quantification was 0.9729. We identified protein changes in malignant tissues compared to matched control samples in HCC patients, and further stratified patient samples into groups with high α-fetoprotein (AFP) expression or HBV infection. In aggregate, the data identified 23 upregulated pathways and 13 ones. We observed enhanced biomolecule synthesis and suppressed small molecular metabolism in liver tumor tissues. 16 proteins of high documented relevance to HCC are highlighted in our data. We also identified changes of virus-infection-related proteins including PKM, CTPS1 and ALDOB in the HBV^+^ HCC subcohort. In conclusion, we demonstrate the practicality of performing proteomic analysis of biopsy-level tissue samples with PCT-SWATH methodology with moderate effort and within a relatively short timeframe.

## Introduction

Hepatocellular carcinoma (HCC) is the fifth frequent malignancy worldwide and ranks as the third leading cause of cancer-related mortality^1^. In China, HCC accounts for more than 300,000 deaths every year ^2^. In early-stage disease, HCC patients’ survival can be significantly improved by treatment including surgical resection and liver transplant. However, HCC mostly exhibits no symptom at early stages. Consequently, less than one third of newly diagnosed HCC patients are eligible for potential curative therapies. Therefore, early diagnosis of HCC is vital for patient survival, and the identification of biomarkers for early HCC detection is a crucial clinical need^3^.

Currently three FDA approved serum biomarkers are recommended to indicate the risk of liver cancer in high risk populations: α-fetoprotein (AFP), AFP-L3, which is a fucosylated isoform of AFP, and des-gamma-carboxy-prothrombin (DCP) ^4^. However, these markers are not included in the surveillance guidelines published by the American Association for the Study of Liver Diseases ^5^ and the European Association for the Study of the Liver ^6^ because of low sensitivity and specificity. In China, AFP is used jointly with ultrasonography, computed tomography (CT) and pathology examination for early screening of HCC patients, as recommended by the Asian Pacific Association for the Study of the Liver ^7^. This is based on the knowledge that the positive rate of AFP in HCC is about 60-80% ^8^.

Better biomarkers are needed for HCC. The search for HCC biomarkers from clinical specimens using proteomics approaches has been focused mainly on blood samples ^9,10^. Besides, some studies also investigated biological fluids including urine ^11^ and tissue interstitial fluid ^12^. Nevertheless, HCC tissue samples have not been extensively studied by advanced proteomics methods, even though dysregulated tissue proteins are presumably the main source of potential blood and biofluidic biomarkers. From the literature, we found that Li, et al studied malignant and matched non-tumorous tissues (about 100 mg tissue per sample) from eleven HCC patients using two dimensional LC coupled with shotgun proteomics with a LC gradient of 165 min and observed upregulation of SET Complex proteins ^13^. Megger, et al nominated 51 differentially expressed proteins from about 573 proteins identified by 2D gel electrophoresis and label-free shotgun proteomic analysis of 7 pairs of HCC tissue samples, ^14^. In a more recent study, Naboulsi W. *et al.* identified and quantified 2,736 proteins from 19 pairs of HCC tumorous and adjacent non-tumorous tissue using the label-free shotgun proteomics over a 98-min LC gradient ^15^. Gao et al ^16^ applied SWATH-MS to study 14 pairs of HCC and non-HCC tissues and quantified 4,216 proteins in at least one biological replicates and 1,903 proteins in at least three out of five biological replicates using a 120-min LC gradient. For each sample, about 200 mg tissue was used. These studies are generally of relatively low throughput and consumes relatively large amount of tissue samples.

We have recently developed a methodology integrating pressure cycling technology (PCT) and SWATH mass spectrometry (MS) for the rapid acquisition of proteotypes, defined as the acute quantitative state of a proteome, from biopsy-level tissue samples (about 2 mg wet weight) ^17^. PCT technology produces high pressure (up to 45,000 p.s.i.) and lyses tissue samples by oscillating the pressure between low and high values, thereby allowing tissue lysis and protein digestion to occur in the same PCT-MicroTube. The method can be applied in a semi-automated and standardized manner and achieves relatively high sample throughput ^17–19^. SWATH-MS is an emerging data-independent acquisition (DIA) mass spectrometric method that offers a high degree of quantitative accuracy, proteomic coverage, reproducibility of proteome coverage and sample-throughput ^20^. The PCT-SWATH methodology was further demonstrated to be applicable to rapid proteotype acquisition from sub milligrams of tissue samples ^19^. In our first study that demonstrated the technology, the PCT-SWATH method was applied to process and convert 18 biopsy samples from nine patients with renal cell carcinoma into SWATH-MS fragment ion maps ^17^. With a two-hour LC gradient, a 32 fixed SWATH windows setting, and the OpenSWATH data analysis tool ^21^, about 2,000 SwissProt proteins were identified with a high degree of reproducibility across all samples ^17^. More recently, a more advanced and miniaturized device, *i.e.* PCT-MicroPestle ^18^ was introduced providing for higher efficiency of protein extraction and peptide generation from biopsy-level tissues.

In this study, we firstly improved the PCT-SWATH methodology by adopting a 45-min LC gradient and an acquisition method using 67 variable SWATH windows to increase the sample throughput without compromising the proteomic depth. Then we applied the improved workflow to analyze peptide samples from a cohort of HCC patients in technical duplicate with unprecedented reproducibility and throughput.

## Materials and Methods

### Patients and tissue samples

38 fresh frozen tissues from 19 HCC patients were collected from Union Hospital, Tongji Medical College, Huazhong University of Science and Technology, Wuhan, China. The clinical and pathological data for these patients are provided in Table 1. For each patient, two tissue biopsy punches (with dimensions of 5∗5∗5 mm^3^) including a tumorous tissue and a non-tumorous tissue from an adjacent region as determined by histomorphological examination were collected. All 38 tissue samples were collected from hepatectomy specimens within 1 hour after surgical removal, snap frozen and stored at −80°C until proteomic analysis. The sample collection was approved by institutional review board of the Union Hospital in Wuhan, China. This study was approved by Ethics Committee of Tongji Medical College, Huazhong University of Science and Technology.

**Table 1.**
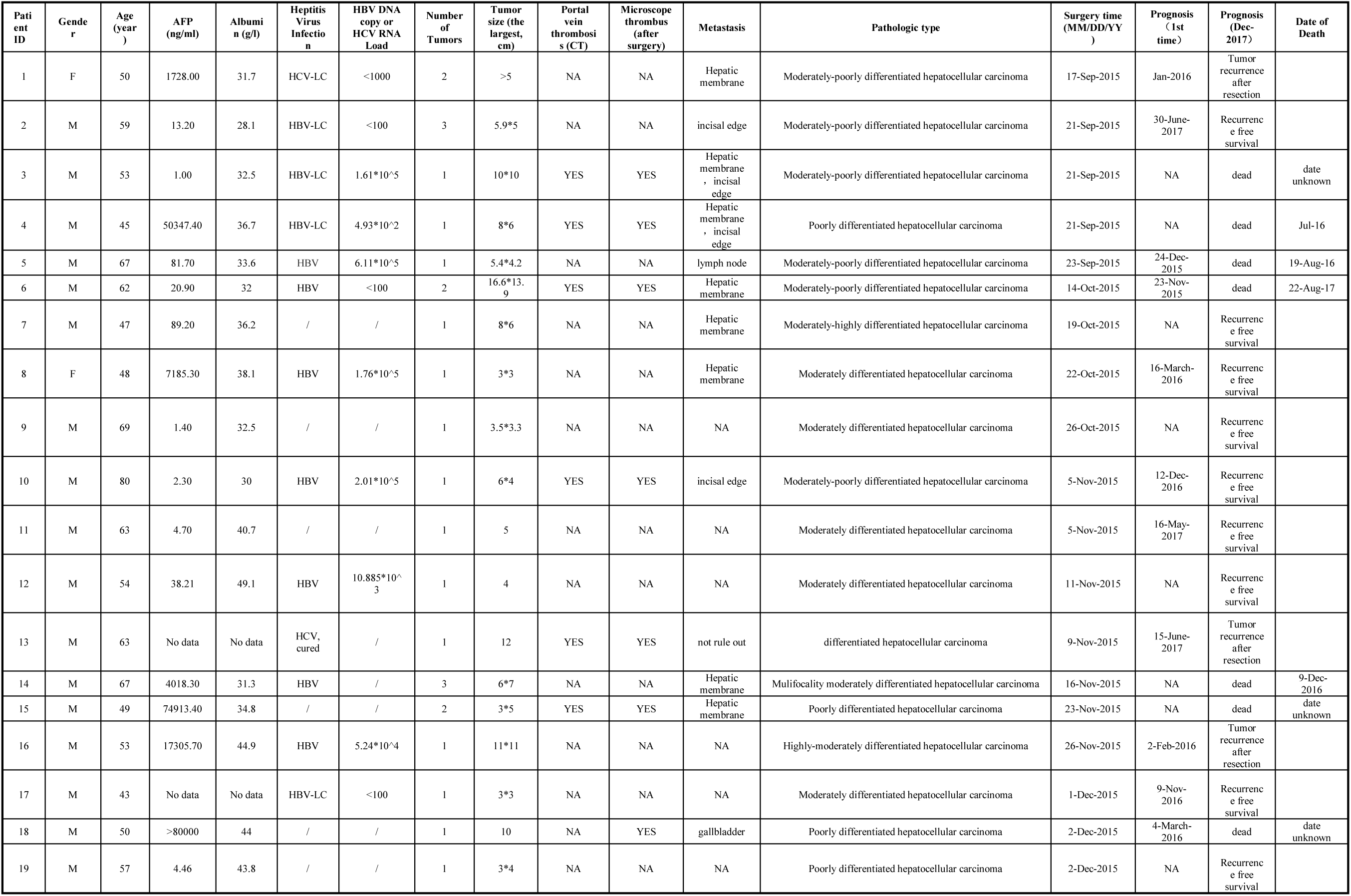
Clinicopathological data of the 19 HCC patients.

### PCT-assisted sample preparation

For each sample, about 1.6 ~ 2.5 mg (wet weight) of frozen tissue was processed based on the protocol described previously ^17,18^. Briefly, tissues were lysed with the PCT-MicroPestle device (Pressure BioSciences Inc, South Easton, MA) in 30 μL of lysis buffer composed of 8M urea, 0.1 M ammonium bicarbonate, and protease inhibitor (Roche) in a barocycler HUB440 (Pressure BioSciences Inc). The tissue lysis was performed under a program consisting of 60 oscillating cycles, with each cycle consisting 50 s of high pressure at 45,000psi and 10 s of ambient pressure at 33 °C. Then the extracted proteins were reduced and alkylated by incubation with 10mM Tris (2-carboxyethyl) phosphine (TCEP) and 20 mM Iodoacetamide (IAA) at 25°C. This was followed by vortexing at 600 rpm for 30 min in the dark, at ambient pressure. Afterwards, digestion was first performed with Lys-C (Wako; enzyme-to-substrate ratio, 1:40) in the barocycler (20,000 psi, 50 s high pressure and 10 s ambient pressure, 45 cycles), followed by trypsin (Promega, enzyme-to-substrate ratio, 1:50) digestion in the barocycler (20,000 psi, 50 s high pressure and 10 s ambient pressure, 90 cycles). After digestion, the peptides were acidified with trifluoroacetic acid (TFA) to pH 2-3 and cleaned with SEP-PAK C18 (1cc 50mg) cartridges (Waters).

### SWATH-MS

The sample samples were spiked with iRT peptides (Biognosis, Zurich, Switzerland) at a 1:10 (vol/vol) ratio. One microgram of peptide sample was injected to an Eksigent 1D+ Nano LC systems (Eksigent, Dublin, CA) and analyzed in a 5600 TripleTOF mass spectrometer (SCIEX) in SWATH mode as described previously ^17^. We optimized the LC gradient to 45 min (Supplementary Fig. 1), and the SWATH acquisition scheme to 67 variable windows (Supplementary Table 1). The other SWATH parameters were set exactly as in our previous study ^17^, except that the ion accumulation time for each SWATH window was 40 ms. Ion accumulation time for peptide precursors was set at 50ms. The 38 samples were injected into the mass spec in randomized sequence once and then the same sequence was injected again to obtain a duplicate analysis. After each gradient, the column was washed twice using ramping gradient to minimize carry-over (Supplementary Fig. 1). Mass calibration using beta-gal was performed every fourth injection.

### OpenSWATH analysis

OpenSWATH (version 2014-12-01-154112) ^22^ was performed as described previously ^17^ except that a pan-human library (32 windows version) ^21^ was used to search the SWATH data in the iPortal platform ^23^. The details of the pan-human library, containing 139,449 peptides from 10,316 proteins, have been described previously^21^. The default parameters as used previously ^17^ are provided in Supplementary Table 2.

### Pathway and process enrichment analysis

Pathway and process enrichment analysis of the differentially abundant proteins were performed using Metascape (http://metascape.org). Terms with p-value < 0.01, minimum count 3, and enrichment factor > 1.5 (enrichment factor is the ratio between observed count and the count expected by chance) are collected and grouped into clusters based on their membership similarities. To further capture the relationship among terms, a subset of enriched terms was selected and rendered as a network plot, where terms with similarity > 0.3 are connected by edges.

Protein-protein interaction enrichment analysis was carried out for each given gene list (http://metascape.org). The resultant network contains the subset of proteins that form physical interactions with at least another list member. Molecular Complex Detection (MCODE) algorithm ^24^ was applied to identify densely connected network components. Pathway and biological process enrichment analysis was applied to each MCODE components independently and the three best-scoring (by p-value) terms were retained as the functional description of the corresponding components.

### Statistics

In volcano plots, paired Student t test was used to compute p values followed by Benjamini & Hochberg correction in R.

### Data deposition

All the SWATH Data files, library and original OpenSWATH results are deposited in PRIDE project ^25^: PXD004873. Reviewer account details: Username; Password: WTpBL22V

*(Note for reviewers: Prior to publication, this data set only be found in PRIDE after the reviewer logs into PRIDE using the username and password provided above. This data set cannot be found without logging in.)*

## Results and Discussion

### Identification of peptides and proteins by PCT-SWATH

To start we extracted proteins from 38 biopsy-level liver tissue samples and produced tryptic peptides using the PCT method ^17,26^. We measured the wet weight of each tissue sample and the total amount of peptide mass generated from the tissue and computed the peptide yield in weight percent per milligram of tissue. The results shown in Fig. 1A indicate that the peptide yield for the 38 samples varied from 1.0% to 9.5%, with an average value of 6.0%. 59.6 μg of total peptide mass per milligram wet liver tissue were obtained on average. The peptide yield in tumor tissue (4.9% on average) was lower than that of normal tissue (8.1% on average, *p* value = 0.00037) (Fig. 1A). The results further showed that tumorous samples exhibited more variability in terms of peptide yield compared to the normal tissue samples. The coefficient of variation (CV) of peptide yield was 15.8% for normal tissue samples (n = 19), 40.7% for tumorous samples (n = 19), and 32.9% overall (n = 38) (Fig. 1B), demonstrating the heterogeneity of liver tumors.

**Figure 1.**
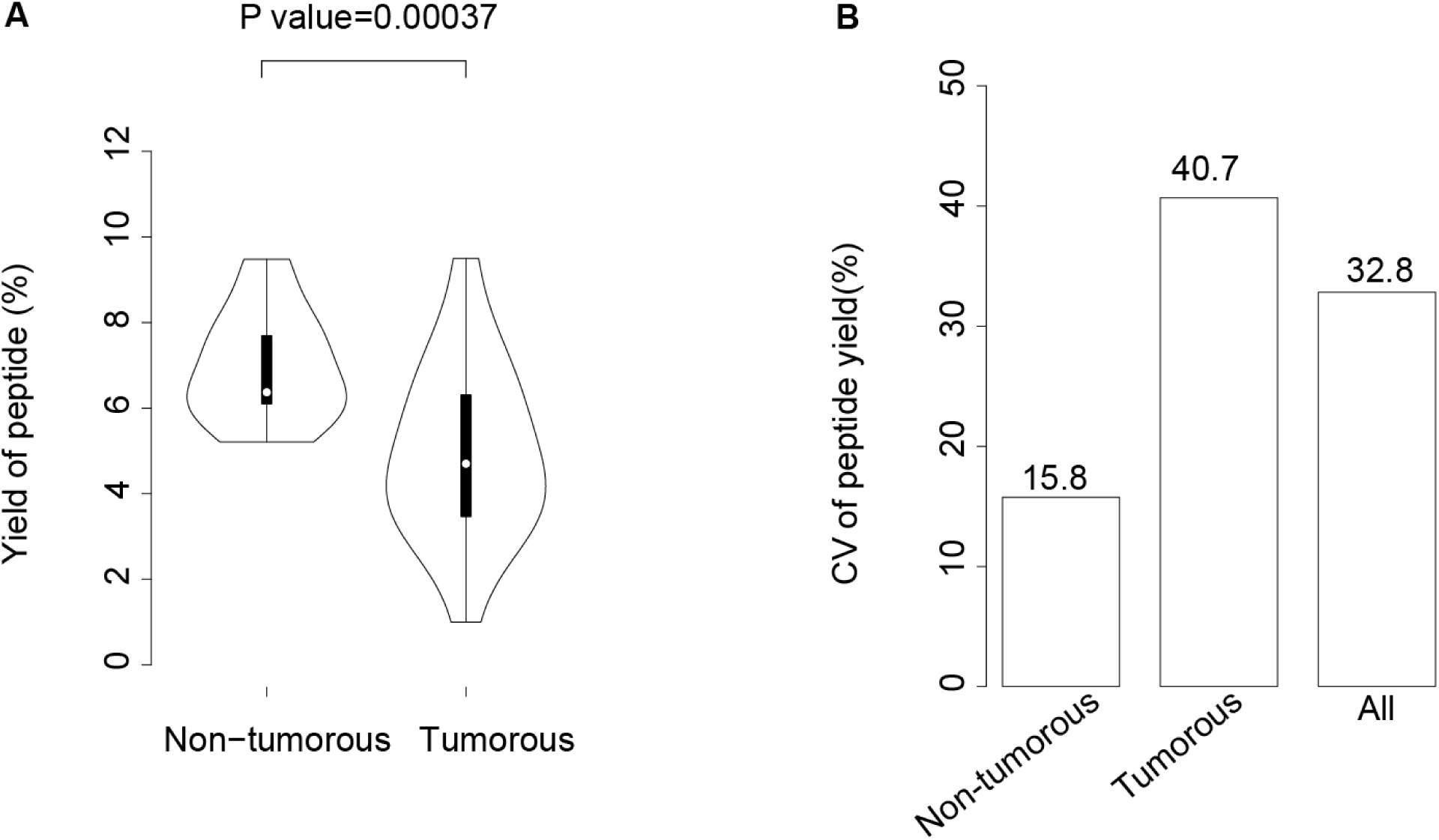
Performance of PCT-assisted peptide preparation. (**A**) Yield of peptide in two types of liver tissue samples. Y-axis shows the peptide yield in percentage of peptide (μg) per milligram (mg) tissue. (**B**) Coefficient of variation (CV) of peptide yield in two types of tissues and all samples.

Each peptide sample was then analyzed using SWATH-MS in technical duplicate, using a 45-min LC gradient and a 67 variable SWATH window acquisition scheme (Supplementary Table 1) which was optimized based on the ion density map of human proteins extracted from HCC tissues. With this SWATH window setting, up to 20 samples were analyzed per MS instrument per 24 hours. Mass calibration using beta-gal was performed every fourth injection. The MS analysis of the 76 SWATH runs were completed in about 4 working days.

After searching the resulting raw data files against the pan-human SWATH assay library ^21^, 2,579 unique proteotypic SwissProt proteins with 11,787 unique peptides were identified at a false discovery rate (FDR) below 1% using OpenSWATH ^21^ (Supplementary Table 3). Comparable numbers of proteins and peptides were quantified from both non-tumorous and tumorous type of tissue samples (Fig. 2A and 2B). In this study, the PCT-SWATH throughput was further increased compared to the previous work ^17^. The protein numbers identified per sample was slightly higher than those obtained from human kidney tissues ^17^.

**Figure 2.**
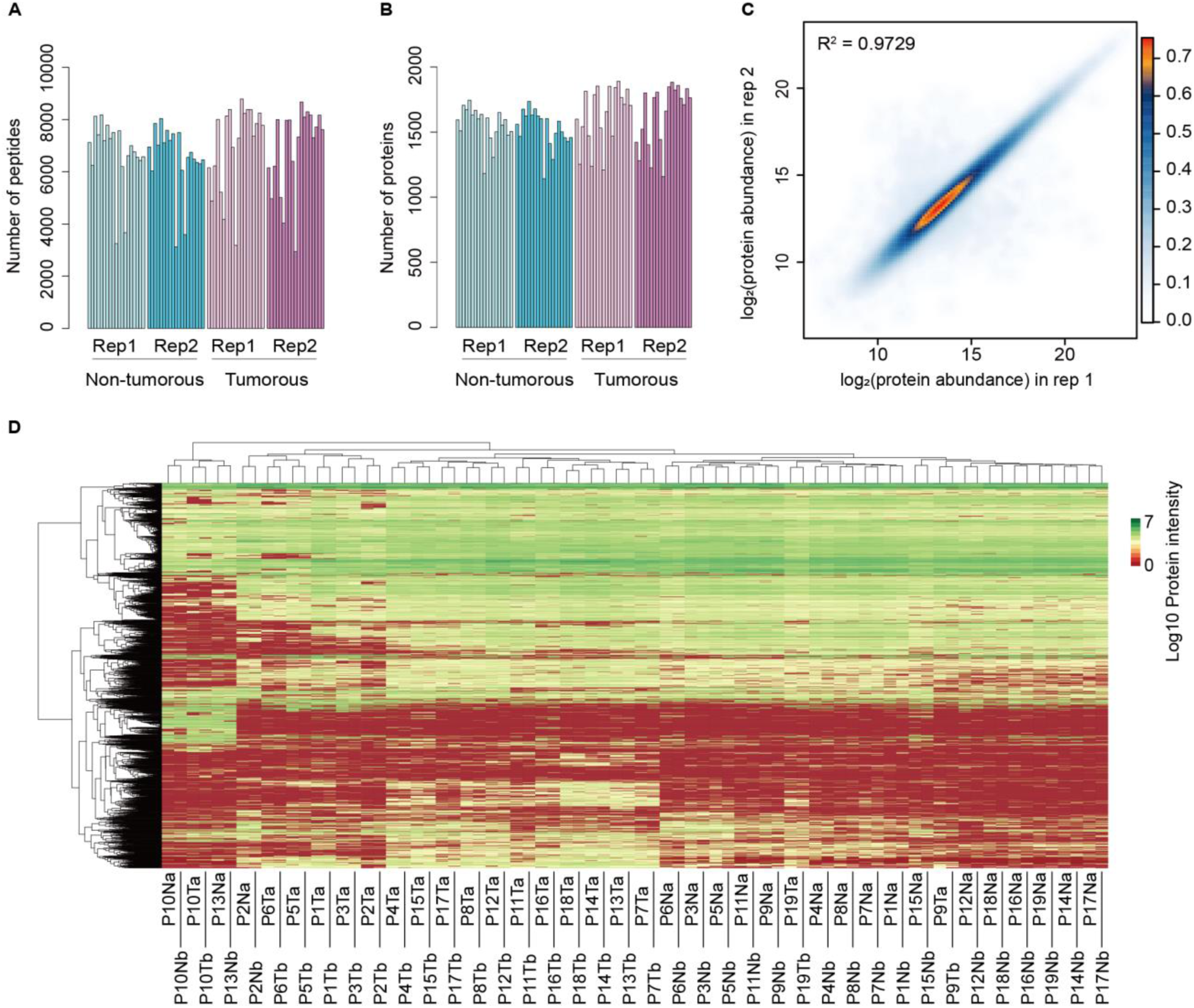
Peptide and protein identification and quantification. (A) Number of peptides identified in each sample (76 SWATH runs from 38 of samples from 19 pairs of HCC patient tissue). (B) Number of proteins identified in each sample. (C) Global Pearson correlation of technical replicates of peptide quantification for samples in all samples. Values are log2 transformed protein intensity. Missing values were excluded from this analysis. (D) Heatmap overview of protein abundance patterns in 76 samples. Protein intensity values were log10 transformed and subjected to unsupervised-clustering in two axes. (P, patient ID; N, non-tumorous; T, tumorous; a, the first technical replicate; b, the second technical replicate; for example, P1Na, the technical replicate run No. 1 of non-tumorous tissue from patient No. 1).

### Variability of protein expression between tumor and non-tumorous tissues

To evaluate the quantitative accuracy of the method, we computed the global Pearson correlation of the two technical replicates at the peptide level for all samples after removing missing values. The R^2^ was 0.9729 (Fig. 2C), indicating that our label-free method could quantify peptides with high degree of reproducibility. The global view of protein quantification is shown in Fig. 2D. It shows that all the technical replicates were clustered together respectively.

We then analyzed the overall variation of protein abundance for all 2,579 quantified proteins from tumor tissues, adjacent non-tumorous (normal) tissues and all samples, respectively (Fig. 3A). The data showed that the CV of tumorous tissue samples was slightly greater than that of adjacent non-tumorous tissues, indicating more overall variability of protein expression in tumors. We next checked the relationship between protein abundance and the overall variation among these samples and observed that low abundance proteins exhibited a higher degree of overall variability (Fig. 3B, 3C, 3D). In Fig. 2C, proteins of low abundance demonstrated comparable reproducibility with high abundance proteins, therefore we mainly attributed the higher variability in tumor tissue to biological causes.

**Figure 3.**
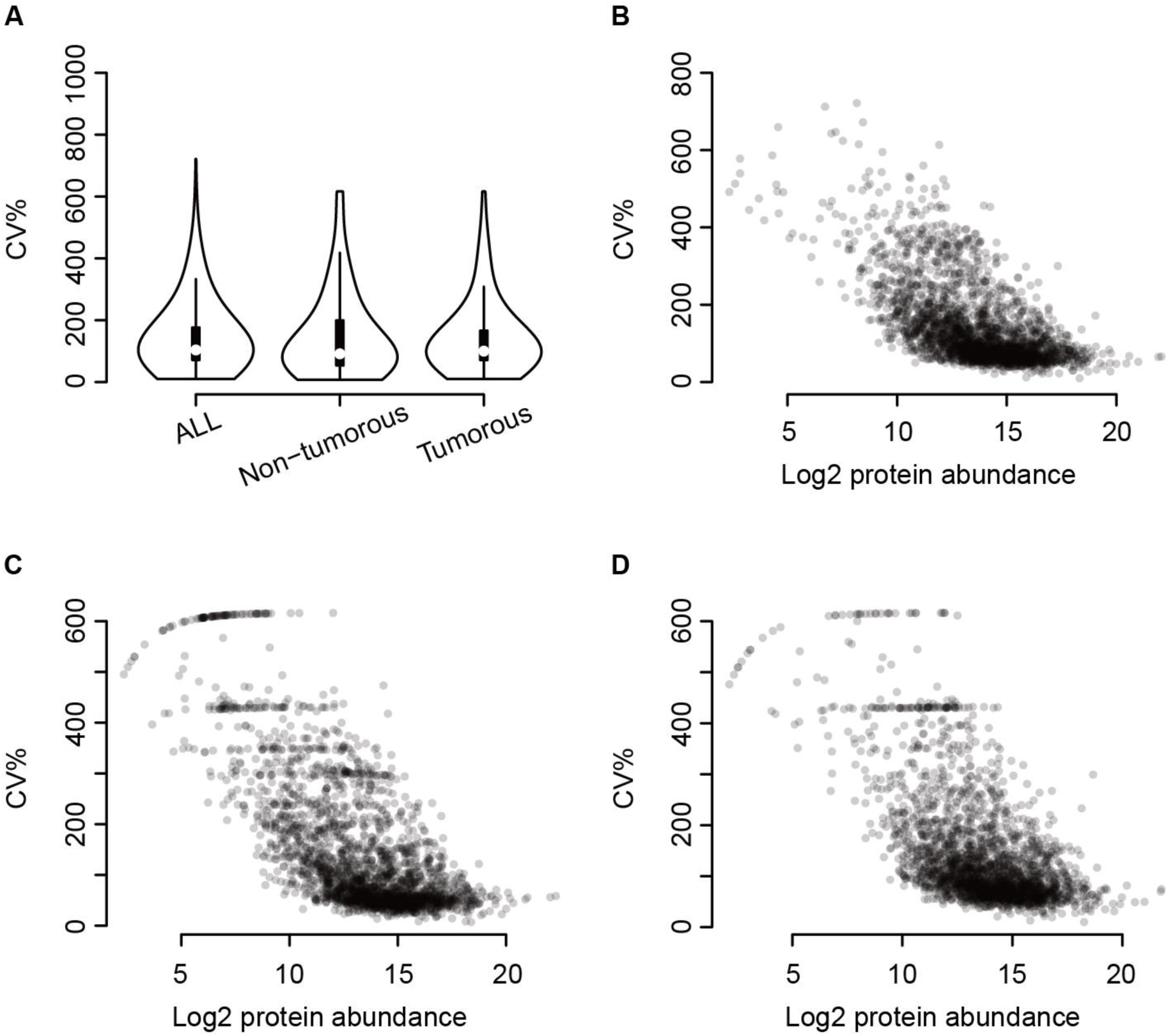
Variability of protein expression in liver tissues. (A) Distribution of protein abundance variability, quantified as percentage of CV values, in tissue samples. B, C and D show correlation of CV of protein abundance variability and protein abundance of all samples, non-tumorous samples and tumorous samples, respectively.

### Differentially regulated proteins characterized in the HCC cohort

We further explored differentially expressed proteins between tumorous and non-tumorous samples. A total of 541 proteins showed significant differential abundance (Fig. 4A). Of these, 381 proteins showed increased abundance and 160 proteins showed decreased abundance in tumorous tissues compared to non-tumorous tissues. Pathway enrichment analysis of the regulated proteins in HCC tumorous tissues was performed by Metascape. We found that proteins with increased abundance were highly enriched in mRNA related processes including mRNA metabolic process, peptide metabolic process, ribonucleoprotein complex biogenesis, and nuclear transport. The proteins with decreased abundance were found to be mostly involved in small molecule metabolic processes such as small molecule catabolic process, monocarboxylic acid metabolic process, and carbon metabolism.

**Figure 4.**
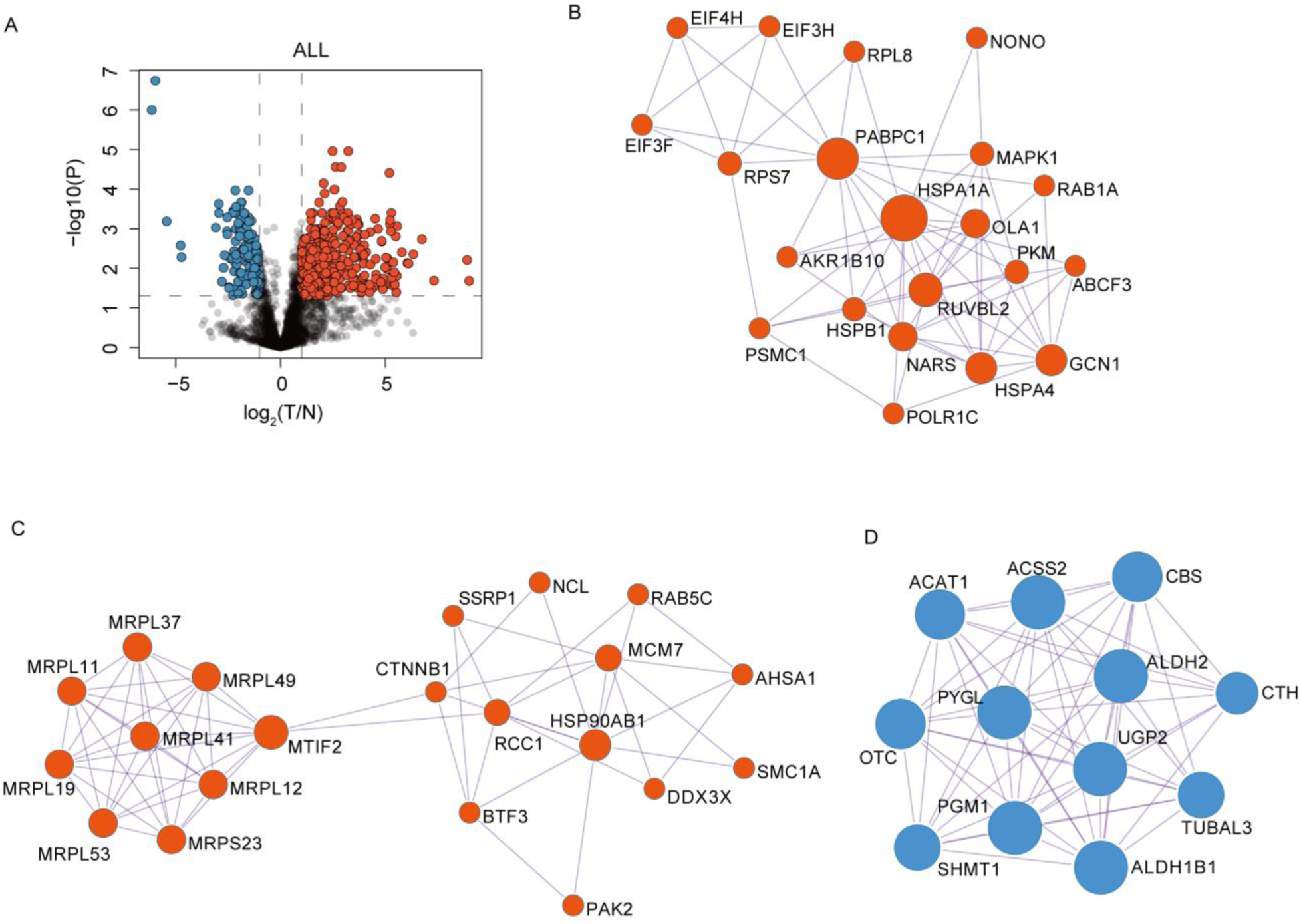
Significantly regulated proteins in all tumorous tissues versus non-tumorous tissues. (A) Volcano plot of proteins expressed in tumorous tissue samples and their matched non-tumorous samples. Each data point represents the ratio of tumorous to adjacent non-tumorous tissue observed for a protein. The horizontal line at –log10 (p-value) = 1.3 indicates p= 0.05. Data points above the line (p < 0.05) are considered to be significantly regulated. The two vertical lines at log2 ratio (tumorous to non-tumorous) = ±1 where the fold change is equivalent to 2 indicate the threshold for up-/down-regulation. The upper left quadrant (p < 0.05 and fold change <0.5) represent the downregulated proteins whereas the upper right quadrant indicates the upregulated proteins in tumorous tissue. N, non-tumorous; T, tumorous. (B) and (C) enriched significantly upregulated protein networks. (D) enriched significantly downregulated protein network.

Next, we carried out protein-protein interaction network analysis using Metascape. The resultant network contains the subset of proteins that form physical interactions with at least another protein showing altered abundance. Heat shock proteins (HSP) were found to be increased (Fig. 4B). HSP proteins constitute a family of highly conserved stress response proteins that can protect cells and induce them to repair damage caused by a variety of stress ^27^. Several members of the HSP family have been reported to be overexpressed in a wide range of human cancers ^28^. The HSP70 family contains at least eight homologous chaperone proteins that differ from each other by amino acid sequence, expression level and sub-cellular localization ^29^. Hsp70-1 is an essential molecular chaperon upregulated in response to heat, stress, or cell survival protection and its function is tightly controlled by heat shock factor-1 (HSF-1). It has been demonstrated that 14-3-3σ protein induces HSP70 via a β-catenin/HSF-1-dependent pathway, which consequentially modulates HCC. The HSF-1/Hsp70-1 protein complex is therefore considered to be targeted for developing a prognostic tool for HCC ^30^.

We also observed that proteins related to mitochondrial translation, specifically the mitochondrial ribosomal proteins (MRPs), were among the most strongly upregulated protein clusters (Fig. 4C). Tumor initiation and progression in cancer cells involve the development of mechanisms to inhibit apoptosis at multiple stages, whereas mitochondria play crucial roles in the induction of apoptosis and programmed cell death. This paradigm is central to malignant cellular transformation because altered mitochondrial function and defective apoptosis are well-known hallmarks of cancer cells. It has also been reported that the expression of genes encoding MRPs, mitoribosome assembly factors and mitochondrial translation factors is modulated in multiple cancers, which is linked to tumorigenesis and metastasis ^31^. Mini-chromosome maintenance complex component 7 (MCM7) was also found to be overexpressed (Fig. 4C). MCM7 plays an essential role in initiating DNA replication. DNA replication is a central process in cell proliferation, while aberrant DNA replication is a driving force of oncogenesis. It has been reported that overexpressed MCM7 is associated with a poor prognosis of HCC patients, and MCM7 promotes HCC cell proliferation via upregulating MAPK-cyclin D1 pathway both *in vitro* and *in vivo* ^32^. In addition, the hallmark MYC targets proteins, and the cell cycle checkpoint proteins involved in G2/M transitions, are also among the upregulated protein networks.

Small molecule catabolic processes define the group of proteins that showed the strongest reduction of abundance in the HCC samples tested (Fig. 4D). Our data showed that proteins involved in glycogen metabolism pathway were suppressed in HCC samples. These proteins include the glycogen synthesis enzymes including phosphoglucomutase-1 (PGM1), and UTP-glucose-1-phosphate uridylyltransferase (UGP2), as well as the glycogen degradation enzyme liver glycogen phosphorylase (PYGL). They are downregulated in tumorous tissues compared to their matched non-tumorous controls, indicating the reprogramming of glycogen metabolism in HCC which suppresses the turn-over of glycogen. However, our data did not indicate whether this reprogramming changed the direction of glycogen storage.

Both cystathionine β-synthase (CBS) and cystathionine-γ-lyase (CTH) were found at decreased abundance in the HCC samples tested (Fig. 4D). CBS and CTH are abundant proteins in the liver, endogenously produce hydrogen sulfide (H_2_S), a gasotransmitter modulating synaptic transmission, vasorelaxation, angiogenesis, inflammation, and cellular bioenergetics. Regulation of H_2_S influences lipid metabolism, glucose metabolism, oxidative stress, and mitochondrial bioenergetics ^33^. Suppression of CBS transcripts has been reported in a study of 120 HCC patients compared to non-cancerous liver tissue ^34^ Reduced CBS transcripts expression is significantly correlated with high tumor stage, high Edmondson grade, and high AFP level ^34^. However, neither the CBS nor the CTH protein level has yet been investigated in HCC tissue. Our data suggests downregulation of H2S production via suppressing CBS and CTH in HCC tissues.

The ethanol metabolism was also found to be downregulated (Fig. 4D). Aldehyde dehydrogenase (ALDH) is a gene superfamily that is responsible for the detoxification of biogenic and xenogenic aldehydes. Both ALDH2 and ALDH1B1 proteins were significantly downregulated. The ethanol detoxifying pathway in humans occurs mainly in the liver and is carried out by two enzymatic steps. In the first step ethanol is metabolized quickly by alcohol dehydrogenase (ADH) to generate acetaldehyde, and the latter is then metabolized by the mitochondrial ALDH2 to acetate ^35^. ALDH2 is expressed ubiquitously in all tissues but is most abundant in the liver and found in high amounts in organs that require high mitochondrial oxidative phosphorylation. The downregulation of ALDH in tumorous liver tissue will likely accumulate toxin acetaldehyde and promote tumorigenesis.

In summary, liver is the pivotal organ for molecular synthesis and the metabolic hub of human beings. Our data show that the molecular synthesis related pathways are enhanced, while most major metabolic processes were suppressed in the HCC tissues tested, suggesting a biochemical imbalance in HCC cells.

### Proteins with altered abundance in HCC patients with high serum AFP

Next, we explored the molecular pathogenesis of HCC by associating protein expression with various clinic-pathological characteristics of HCC. AFP is the most widely used tumor biomarker currently available for early detection of HCC, and the widely accepted threshold of serum AFP is 20 ng/mL. It has been reported that at this threshold, serum AFP had a sensitivity of 41-65% and specificity of 80-94% ^36^. We grouped the 19 HCC patients in our cohort into two groups, according to an AFP-cutoff of 20 ng/mL (Table 1), computed the fold change of protein expression between tumorous and non-tumorous tissues, and checked the protein regulation pattern of 11 patients with high AFP level (> 20 ng/ml) and six patients with low AFP level (< 20 ng/ml), respectively. Two patients without serum AFP examination value at surgical operation were excluded from these analyses. Although a considerable proportion of HCC patients in the cohort do not have elevated serum AFP, and serum AFP can increase in patients with diseases other than HCC ^4^, high expression of AFP in serum correlated with high cell proliferation, high angiogenesis and low apoptosis and was associated with poor prognosis of HCC patients ^37^. Indeed, we found in our data set that the low serum AFP level was closely related to higher survival rate of the patients (5 out of 7 patients have recurrence free survival), especially when there was no portal vein thrombosis detected by CT at diagnosis, as is shown in Table 1. On the other hand, in the AFP-high patient group, 5 of 10 patients were dead and 2 patients suffered from tumor recurrence, displaying poorer survival (Table 1). Nevertheless, statistical analysis was not feasible due to small sample size.

With respect to the AFP-high patient group, 419 proteins showed increased abundance and 192 proteins showed decreased abundance (P value < 0.05) in tumorous samples compared non-tumorous samples. Pathway enrichment and network analysis of these differentially expressed proteins revealed similar results with the above 541 regulated proteins from all 19 HCC patients (Fig. 5A). In the AFP-low subgroup, our data showed no protein significantly regulated between the tumorous and non-tumorous tissues, probably due to the small sample size.

**Figure 5.**
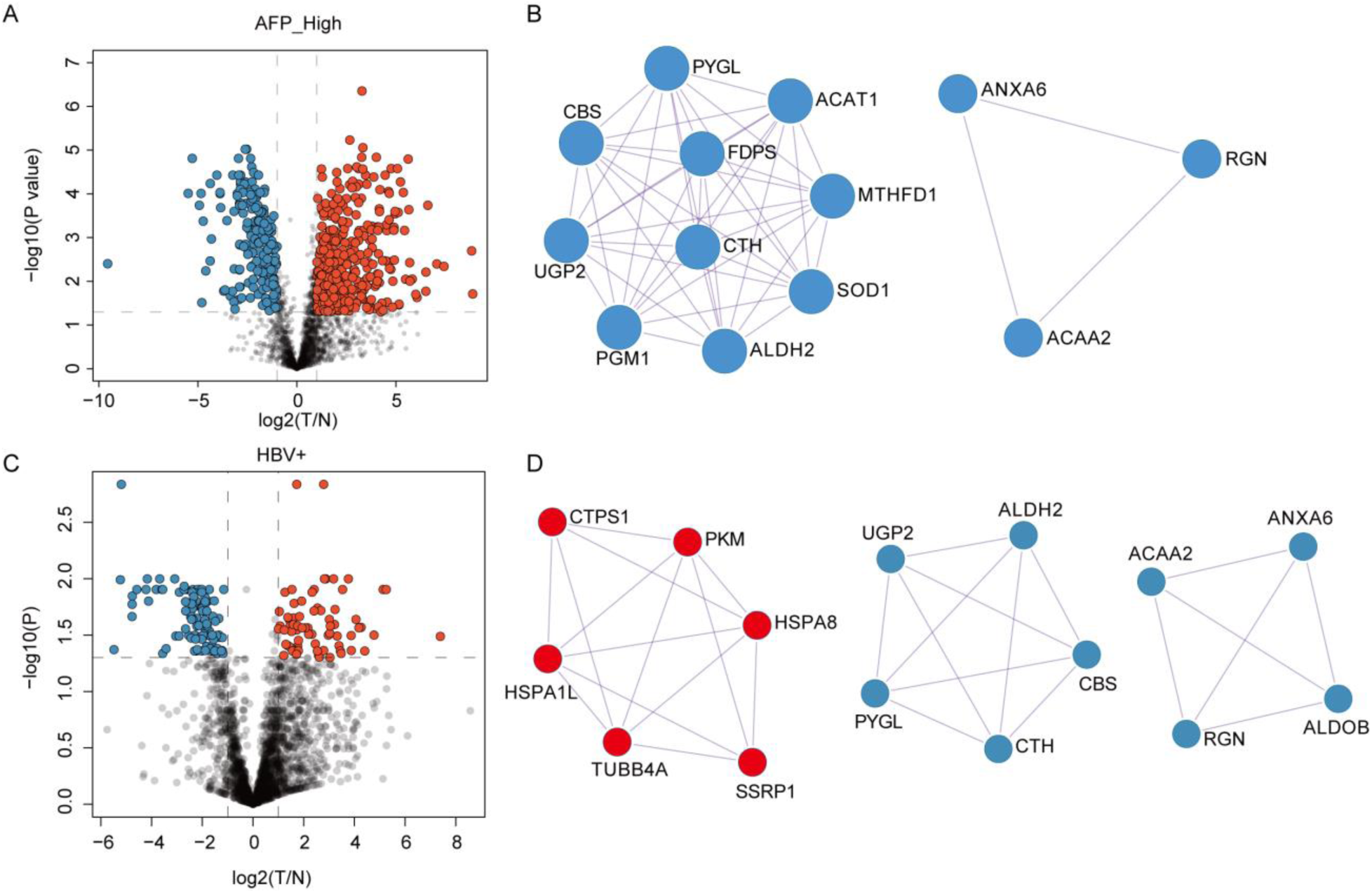
Significantly regulated proteins in tumorous tissues versus non–tumorous tissues grouped by the blood AFP level and HBV infection status. Volcano plots of proteins expressed in tumorous tissue samples and their matched non-tumorous samples in AFP-high patient group (A) and HBV infection status (C). (B) and (D) show the enriched significantly downregulated protein network in the AFP-high patient group and the HBV^+^ group, respectively.

In the AFP-high subgroup (Fig. 5B), we identified additional downregulated proteins including methylene tetrahydrofolate dehydrogenase 1 (MTHFD1), which is an enzyme identified in the one-carbon (1C) metabolism pathway. Another enzyme in the pathway, *i.e.* serine hydroxymethyltransferase (SHMT1), is also downregulated (Fig. 4D). The data suggest reprogramming of one-carbon metabolism in HCC. This is consistent with the Liverome study reporting that downregulation of MTHFD1 is prevalent in proliferative and poorly differentiated HCCs^38^.

We also observed downregulation of a tumor suppressor protein regucalcin (RGN) which modulates multiple protein kinases and phosphatases in cancers including HCC ^39^. Transcripts analysis reports that prolonged survival in HCC patients is associated with increased RGN gene expression ^40^. Our data at protein level further consolidate the potential roles of RGN in HCC.

Two proteins involved in lipid metabolism showed decreased expression in the AFP-high HCC tissues. ACAA2 is a mitochondrial enzyme involved in the fatty acid metabolism pathway. ANXA6 is a lipid-binding protein highly expressed in the liver, regulating cholesterol homeostasis and signaling pathways with a role in liver physiology. A recent study found that ANXA6 was downregulated in HCC tissues compared to non-tumorous tissues in 18 HCC patients ^41^. The data indicate dysfunction of lipid metabolism in HCC tissues.

### Quantitative proteomics analysis of Hepatitis B virus positive (HBV^+^) HCC subgroup

Chronic HBV is the most common cause of HCC in China. Approximately 80% of HCC develops from liver cirrhosis, which predominantly progresses from HBV in China ^42^. We found that 106 and 75 proteins showed increased or decreased abundance, respectively, in 11 HCC patients with HBV infection (Fig. 5C) compared to those without HBV infection. Subsequent pathway and protein interaction analysis by Metascape revealed a number of interesting protein clusters (Fig. 5D) that showed a close relationship to liver cirrhosis in patients with chronic HBV infection. Pyruvate kinase M (PKM) was found to be upregulated in this HBV^+^ subgroup (Fig. 5D). PKM is a protein kinase that catalyzes the final step in glycolysis by transferring the phosphate from phosphoenolpyruvate (PEP) to ADP, thereby generating pyruvate and ATP ^43^. PKM has two isoforms, PKM1 and PKM2. PKM2 plays an important role in metabolic alterations related to inflammation and cancer. The ATP generation by PKM2, unlike mitochondrial respiration, is independent of oxygen and thus allows tumor cells to grow in hypoxic conditions. Cancer cells are characterized by high glycolytic rates to support energy regeneration and anabolic metabolism, along with the high expression of PKM2 ^44^. It has been shown in a recent report that the anti-apoptotic protein poly(ADP-ribose) polymerase (PARP)14 promotes aerobic glycolysis in HCC by maintaining low activity of PKM2, and that the PARP14-JNK1-PKM2 regulatory axis links apoptosis and metabolism ^45^. In agreement with our finding that HBV^+^ tumors expressed higher level of PKM, there is another report showing that high PKM2 expression was more frequently found in cirrhotic liver caused by HBV infection than in non-cirrhotic liver, and that PKM2 overexpression was associated with poor survival rates in HCC patients with cirrhotic liver (CL) ^46^.

Cytidine triphosphate synthase 1 (CTPS1) is another example of a protein that showed higher expression in the HBV^+^ HCC subgroup in this study compared to patients without HBV infection (Fig. 5D). CTPS1 catalyzes the rate-limiting step in the *de novo* CTP synthetic pathway, in which a UTP is converted into CTP with the consumption of glutamine and ATP. In fact, upregulated CTPS1 activity has been observed in multiple human and rodent tumors ^47^, promoting tumor transformation and progression ^48^. Recently, researchers focus on the CTPS cytoophidium that is a filamentous intracellular macrostructure aggregated by CTPS, and is able to promote the activity of CTPS ^49^. A recent study examined the presence of CTPS cytoophidia in the tumorous and the adjacent non-tumorous tissues of HCC patients, and found that many cytoophidia were observed in 28% of the HCC tumor samples but not the adjacent hepatocyte population ^50^. In addition, they found that the high expression of HSP90 was correlated with the presence of CTPS cytoophidia, which is consistent to our findings that heat shock proteins (HSPA1L and HSPA8) and CTPS1 are all upregulated.

Fructose-bisphosphate aldolase B (ALDOB) is an enzyme for glucose and fructose metabolism. In a cohort of 313 HCC patients, ALDOB was found downregulated in HCC tissue using tissue microarray and immunohistochemistry (IHC) technologies, and its downregulation was significantly correlated with HCC progression ^51^. Furthermore, ALDOB has been found to be a binding protein of the S region of the HbsAg ^52^. Our data showed that ALDOB expression suppressed in HBV^+^ HCC patients, suggesting a potentially interesting mechanistic linkage between HCC and HBV infection, which might be utilized as therapeutic target.

### Summary of regulated proteins in HCC samples

We summarize all differentially expressed proteins in HCC samples including the AFP-high group and HBV^+^ group (Fig. 6A). Interestingly, the proteins with increased expression concentrated in three types of pathways: a) production of DNA, mRNA and protein; b) oncogenic signaling pathways and apoptosis; c) immune response. This suggests enhanced proliferation of tumor cells in the malignant tissues. In contrast, proteins participated in various metabolism pathways were down-regulated, indicating the suppression of metabolic functionality in liver during oncogenesis. Compared to a previous report of HCC tissue proteome using SWATH-MS ^16^, our data are acquired in a shorter time frame but identified similar protein alternation. For instance, we also observed malignant tissue displayed increased abundance of glucose-6-phophate dehydrogenase (G6PD) and perilipin 2 (PLIN2), as well as downregulation of formimidoyltransferase cyclodeaminase (FTCD) and phosphoenolpyruvate carboxykinase 2 (PCK2, mitochondrial).

**Figure 6.**
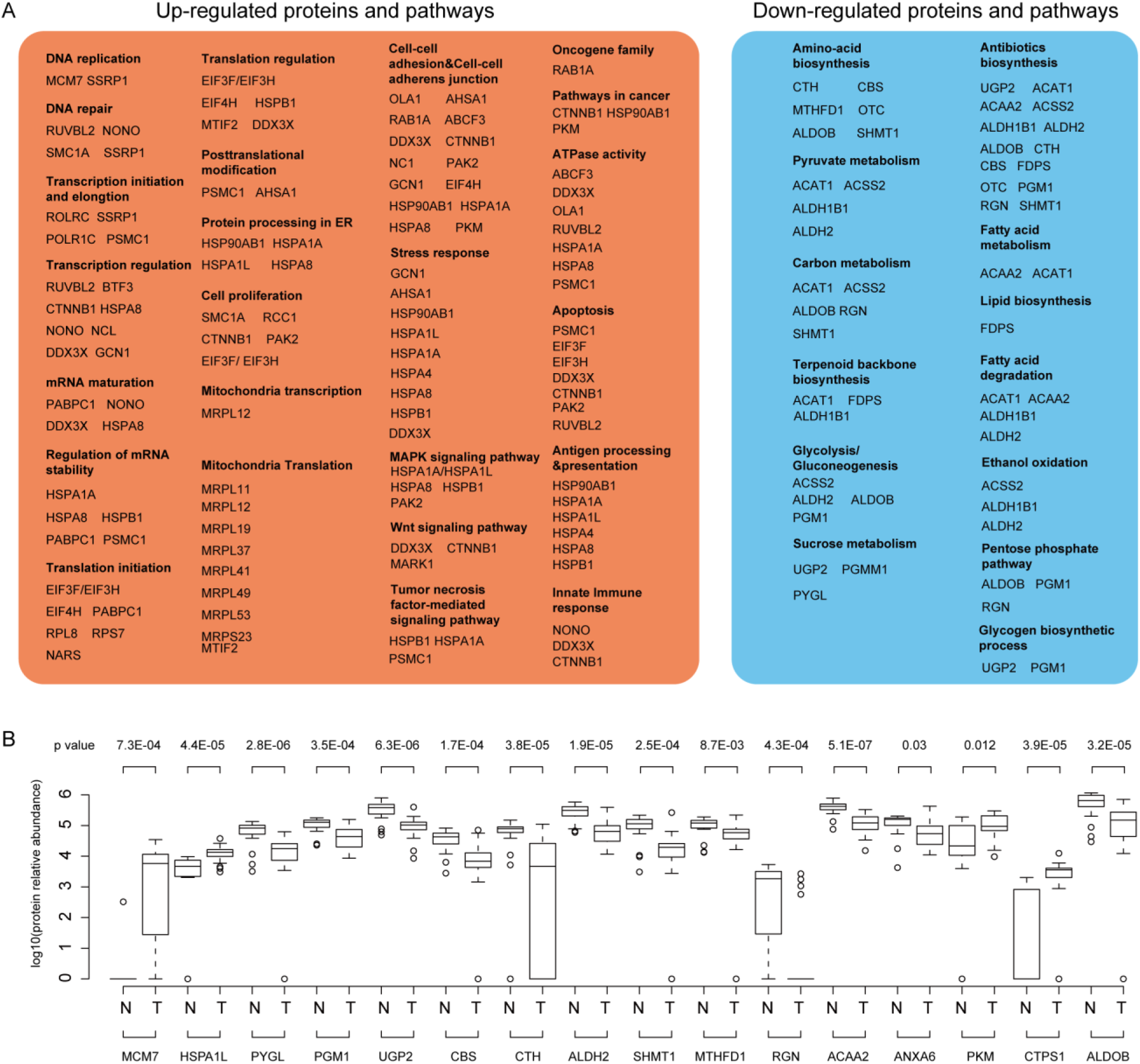
Summary of regulated proteins and pathways in HCC tissues compared to matched control samples. (A) All proteins found regulated in HCC tissues, including AFP-high group and HBV^+^ group, were inspected in literature. Regulated pathways and their component proteins are displayed. (B) 16 selected proteins significantly regulated in HCC tissues compared to matched non-tumorous samples. For each protein, its expression, after 1og10 transformation, in non-tumorous (N) and tumorous (T) tissues are shown in boxplots. Student *t* test was used to compute the significance between N and T samples.

From literature review of all the regulated proteins in Fig. 6A, we identified 16 proteins of clinical relevance to HCC (Fig. 6B). For instance, the overexpression of PKM2 was more frequently found in cirrhotic liver resulted from HBV infection, and ALDOB, whose interaction with HbsAg might be applied as a potential therapeutic target for HBV-related hepatitis or HCC. We calculated the expression level of these 16 proteins between tumorous and the adjacent non-tumorous (named as normal) tissue, as was shown in Fig. 6B, the log_10_ MS intensity, with the *p* value for each pair of comparison between normal and tumor. Although this study is not the first to identify the relevance of these proteins to liver cancer, our study further consolidates their roles in HCC tissues, and more importantly, we demonstrate that this PCT-SWATH-based high-throughput methodology allows measurement of clinically relevant proteins from biopsy-level tissue samples.

## Conclusion

In conclusion, here we report an optimized PCT-SWATH workflow enabling analysis of clinical tissue specimens with unprecedented sample throughput without compromise of quantitative accuracy and proteomic coverage. The entire process (from receiving 38 tissue samples to completing the data acquisition of 76 SWATH files) took about 9 working days, enabling timely measurement in a clinical scenario. Our study reveals a regulated protein landscape in this HCC cohort. Remarkably, proteins with increased abundance are mainly related to mass production, oncogenic signaling and immunity, where metabolic proteins show lower expression. We identified 16 dysregulated proteins of high clinical relevance to HCC, as reported by the literature and substantiated by our data. The study demonstrates that the PCT-SWATH methodology has potential to be practically applied in clinical research to generate high quality proteomic datasets and measure clinically relevant proteins from biopsy-level tissue samples.

## Author contributions

T.G. conceived and coordinated the project. J.Z., C.L., W.X., P.S. performed the PCT-based sample preparation. J.Z., Y.Z., Q.Z., T.Z., T.G. performed the data analysis. Y.Z., J.Z., R.A., S.H., T.G. wrote the manuscript with inputs from all co-authors. R.A., S.H. and T.G. supervised the project.

## Acknowledgements

This study was supported the National Natural Science Foundation, P.R China (NNSF/81200348/2013), the Swiss National Science Foundation (SNF 166435 MitoModules to R.A) ERC PROTEOMICS4D (project-no 670821), start-up package from Westlake Institute for Advanced Study to T.G.. Y. Z. was supported by a fellowship from SignalX (SystemsX RTD 2013/156) and Eu PrECISE (project-no 668858).

